# Global-scale microbiome analyses identify an ancestral gut cyanobacterial lineage enriched in African populations

**DOI:** 10.64898/2026.04.24.720558

**Authors:** Kotaro Murai, Yutaka Suzuki, Suguru Nishijima

## Abstract

Industrialization has profoundly reshaped the structure and function of the human gut microbiome, yet our current knowledge of its diversity remains heavily biased toward industrialized populations. Here, we performed an integrated analysis of a large-scale metagenomic dataset (n = 41,471) to systematically characterize the gut microbiome across the African continent. We show that sub-Saharan African populations harbor gut microbiome structures that are distinct not only from industrialized populations but also from other non-industrialized populations. Notably, we identify a strong enrichment of *Gastranaerophilales*, a poorly characterized lineage of non-photosynthetic Cyanobacteria, in sub-Saharan African populations. Comparative genomic analyses reveal that this lineage encodes distinctive functional features, including motility-related genes, vitamin biosynthesis pathways, and specialized carbohydrate transport systems. Furthermore, we show that this lineage is widely distributed across non-human primates, supporting an evolutionarily conserved host association predating modern humans. Together, our findings identify *Gastranaerophilales* as a potential ancestral gut symbiont that has been markedly reduced in industrialized societies but retained within African populations, highlighting a major but previously underappreciated component of the human gut microbiome.

## Introduction

The human gut microbiome plays a crucial role in maintaining immune and metabolic homeostasis, and its disruption is associated with diverse diseases^1–3^. Recent advances in high-throughput analysis techniques have greatly accelerated microbiome research^4,5^. However, the majority of these studies have focused on populations from industrialized societies, primarily in Europe and North America^6,7^. As a result, current knowledge of the structure and diversity of the human gut microbiome remains incomplete and biased.

Addressing this bias requires the inclusion of populations that retain diverse ecological and lifestyle contexts. Africa is particularly important in this regard. As the origin of modern humans, the African continent harbors the highest levels of human genetic diversity and encompasses a wide range of environments, cultural traditions, and lifestyles that predate industrialization^8^. Previous studies of specific African populations/regions have consistently reported gut microbiome configurations that differ markedly from those of industrialized populations^9–13^. For example, studies of the Hadza hunter-gatherers of Tanzania and other African groups have revealed higher gut microbial diversity and the presence of microbial taxa that are rare or absent in Western populations^14–18^, often referred to as disappearing microbiota or VANISH taxa^19,20^. More recently, the large-scale AWI-Gen 2 Microbiome Project, which analyzed approximately 1,800 adults across four African countries, revealed extensive previously uncharacterized microbial diversity and numerous microbial genomes not captured in previous datasets^21^. These findings highlight the importance of the African gut microbiome for a deeper understanding of human gut microbiome diversity and its association with disease. However, existing studies of African populations have largely focused on specific regions or individual groups, and a comprehensive continent-wide analysis that integrates geographic, cultural, and lifestyle diversity remains lacking.

In this study, we integrate extensive metagenomic and 16S rRNA gene datasets to systematically characterize the structure and function of the gut microbiome in sub-Saharan African populations. This analysis reveals distinct microbiome profiles and uncovers previously underrepresented cyanobacterial lineages largely overlooked in studies of industrialized populations, providing new insights into the evolutionary context of the human gut microbiome.

## Results

### Global diversity and structure of the gut microbiome

We collected and analyzed a large-scale human gut metagenomic dataset comprising 41,471 samples from 57 countries, primarily obtained from the Metalog database^22^ (Methods, Fig. 1a). This dataset spanned a wide range of geographic regions, lifestyles, and socioeconomic backgrounds, including industrialized nations—primarily Western countries and East Asia—as well as diverse populations from non-industrialized countries across multiple continents (Supplementary Tables 1 and 2), thereby providing a comprehensive global representation of the human gut microbiome.

**Fig. 1.**
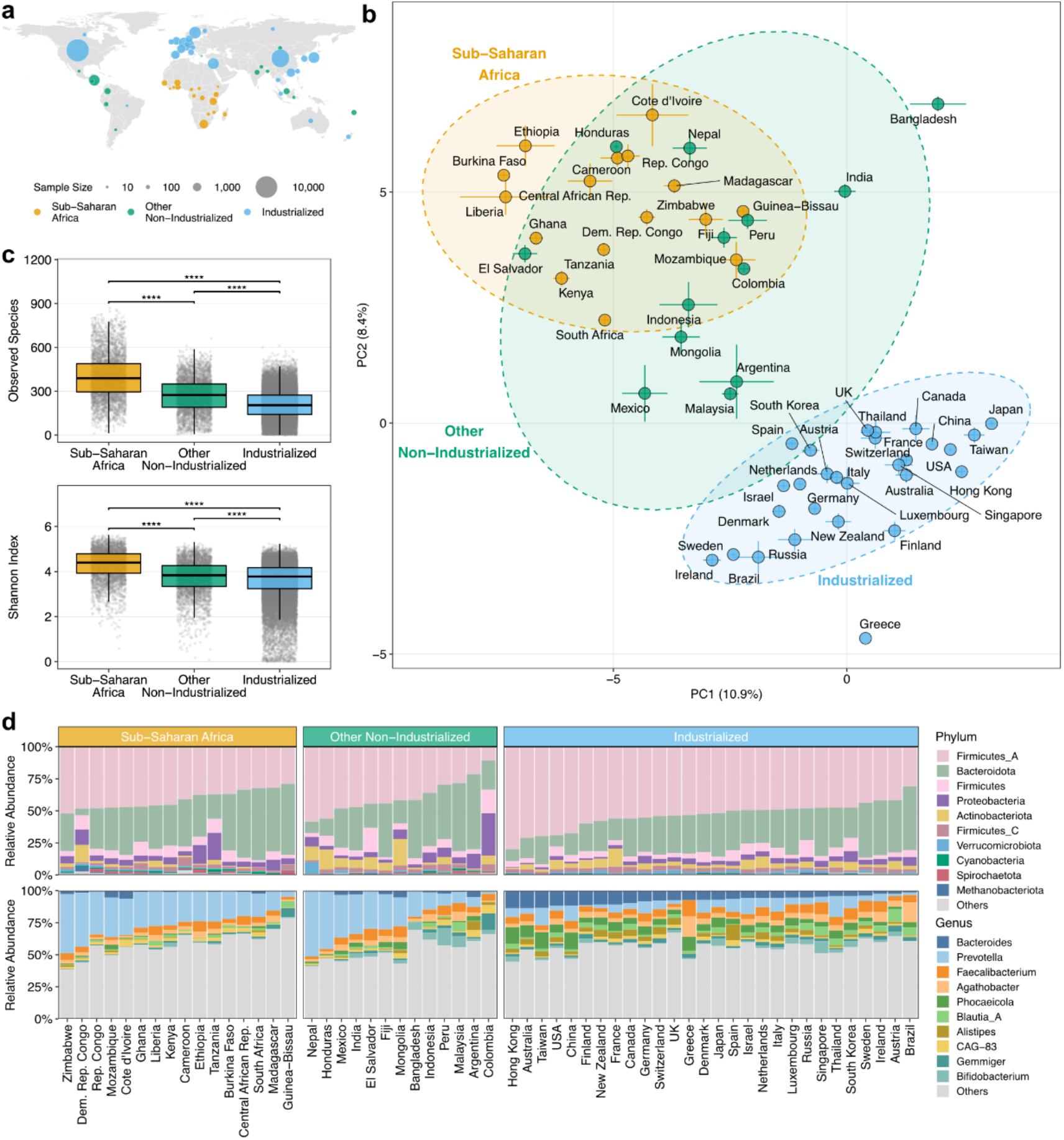
Global landscape of the human gut microbiome based on large-scale metagenomic datasets. **a**, Geographical distribution of the human gut metagenomic samples used in this study (n = 41,471). Circle size indicates the sample size for each country. **b**, Principal component analysis based on species-level relative abundances of the microbiomes. Each point represents the centroid of a country, and error bars denote the standard error of the mean. Countries were classified into three analysis groups (sub-Saharan Africa, other non-industrialized, and industrialized). **c**, Alpha diversity (Shannon index and observed species richness) of the gut microbiome across the three analysis groups. In the box plots, the center line indicates the median, box limits indicate the upper and lower quartiles, and whiskers extend to 1.5× the interquartile range, with individual data points overlaid. (Pairwise comparisons using the Wilcoxon rank-sum test with Bonferroni correction; ****: *P* < 0.0001). **d**, Taxonomic composition of the microbiomes across countries at the phylum (top) and genus (bottom) levels. Stacked bar charts show the average relative abundance of the taxa for each country, aggregated by analysis group. Taxa outside the top 10 most abundant are grouped as “Others”.

Using the species-level taxonomic profiles, we performed a principal component analysis (PCA) to examine global patterns of the gut microbiome variation across countries. The analysis revealed distinct clusters along the first (PC1) and second (PC2) principal components, separating industrialized populations from non-industrialized populations (Fig. 1b). Based on these results, we categorized the countries into three groups reflecting both industrialization and geographic context: “Industrialized”, “Other Non-industrialized”, and “Sub-Saharan Africa”. This stratification enabled us to distinguish African populations from other non-industrialized populations that share similar economic status but differ in ecological and lifestyle contexts. Comparative analysis of alpha diversity revealed a stepwise increase across these groups (Fig. 1c). Both observed species richness and Shannon diversity were lowest in the industrialized group, intermediate in the other non-industrialized group, and highest in the sub-Saharan Africa group, highlighting the exceptional diversity of African gut microbiomes.

At the phylum-level, major phyla such as Firmicutes_A and Bacteroidota were consistently dominant across all three groups (Fig. 1d). At the genus-level profile, *Bacteroides* predominated in the industrialized populations, whereas *Prevotella* was enriched in both non-industrialized groups, irrespective of geographic origin. This shift between *Bacteroides*- and *Prevotella*-dominated microbiomes in industrialized and non-industrialized groups is consistent with previous studies, and has been linked to dietary patterns, particularly the contrast between low-fiber, animal-based diets and high-fiber, plant-based diets^9,10^.

### Systematic identification of gut microbial species characteristic of sub-Saharan African populations

To systematically identify microbial taxa associated with sub-Saharan African populations, we applied linear mixed models (LMMs) that incorporated country and study as random effects to account for geographic and cohort-level heterogeneity (Methods). Compared to both the other non-industrialized and the industrialized groups, we identified 349 species that were significantly enriched and 11 species that were significantly depleted in the sub-Saharan African group (FDR < 0.05, |Estimate| > 0.1, Fig. 2a, Supplementary Table 3).

**Fig. 2.**
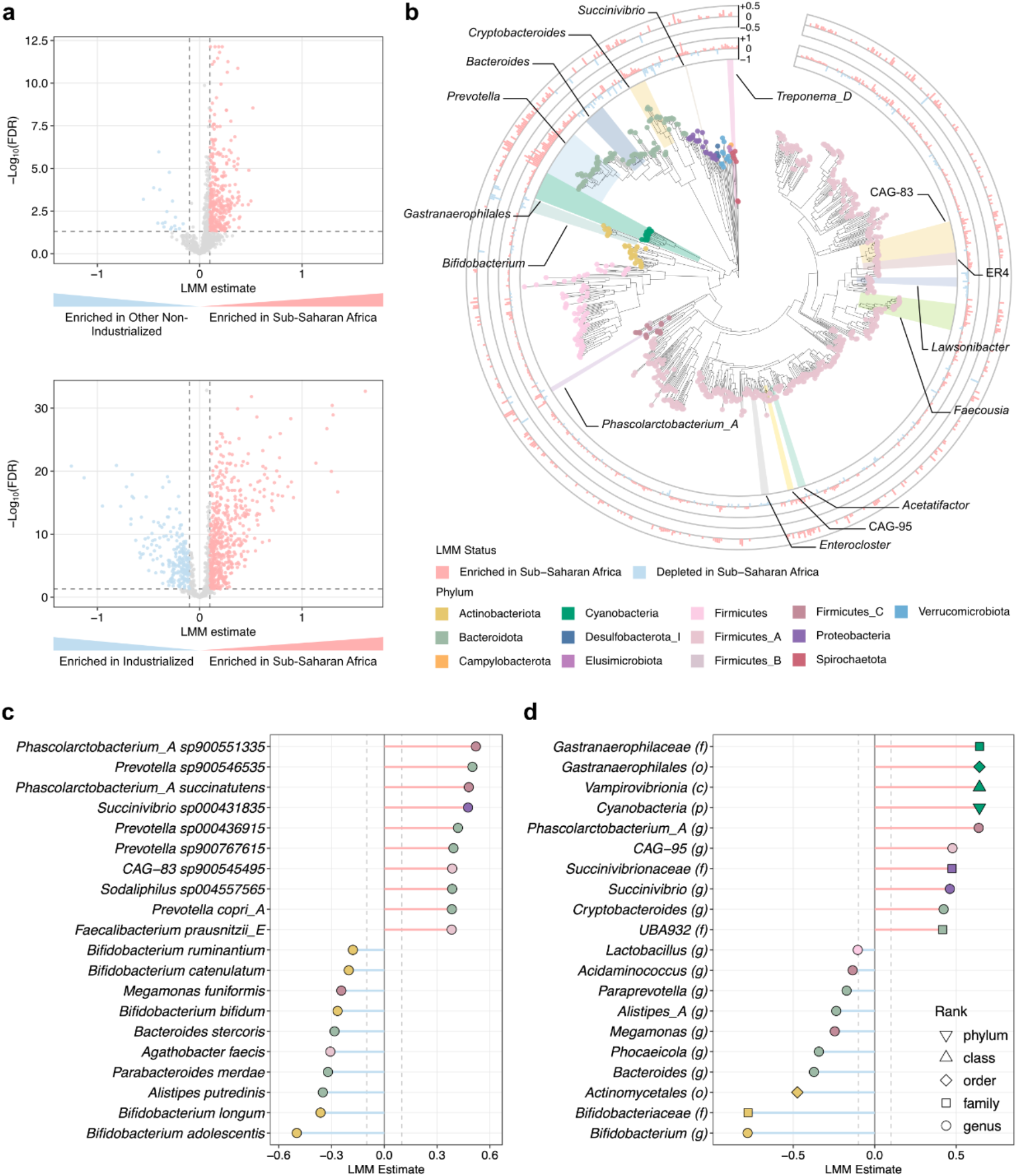
Systematic identification of gut bacterial taxa specific to sub-Saharan African populations. **a**, Volcano plots showing the differential abundance of gut bacterial species in the sub-Saharan Africa group compared with the other non-industrialized group (top) and the industrialized group (bottom). The x-axis represents the linear mixed model estimate based on log_10_-transformed relative abundances (with a pseudocount of 10^−4^). Positive values indicate enrichment in the sub-Saharan Africa group. The y-axis shows the −log_10_-transformed false discovery rate corrected by the Benjamini-Hochberg method. Red and blue points represent species significantly enriched (FDR < 0.05, Estimate > 0.1) and depleted (FDR < 0.05, Estimate < −0.1) in the sub-Saharan Africa group, respectively. **b**, Phylogenetic tree of the analyzed gut bacterial species. The color of the tree tips denotes the phylum-level classification. The two surrounding bar charts illustrate the representative linear mixed model estimates for species exhibiting significant differences. The inner and outer rings represent the comparisons of the sub-Saharan Africa group against the industrialized and other non-industrialized groups, respectively. Bar lengths correspond to the magnitude of these estimates, with red and blue bars representing species significantly enriched and depleted in the sub-Saharan Africa group, respectively. **c**, Representative species significantly altered in the sub-Saharan Africa group (top 10 species with the largest absolute estimates for both enrichment and depletion). The x-axis indicates the LMM estimate (all FDR < 0.05). Point colors correspond to the phylum-level classification in **b. d**, Representative taxa showing significant variation from the phylum to the genus level (top 10 signatures with the largest absolute estimates for both enrichment and depletion). The x-axis indicates the LMM estimate (all FDR < 0.05). Point colors follow the phylum classification in **b**, and shapes denote the taxonomic rank.

The most strongly enriched species in the sub-Saharan African populations included *Phascolarctobacterium*_A sp900551335, *Prevotella* sp900546535, *Phascolarctobacterium_*A *succinatutens, Succinivibrio* sp000431835, and *Prevotella* sp000436915 (Figs. 2b and c). Several *Prevotella* spp. remained significantly enriched in African populations, even compared with the other non-industrialized groups (Supplementary Table 3). We further found that a substantial fraction of enriched taxa in sub-Saharan Africa corresponded to poorly characterized or uncultured species lacking detailed taxonomic annotation, such as CAG-196 sp900553895, CAG-724 sp905202255, and UMGS363 sp900543105. Overall, 49.3% (172/349) of significantly enriched species lacked curated genus-level annotations and were represented by placeholder identifiers (e.g., CAG-, UMGS-, UBA-) in Genome Taxonomy Database^23^. This indicates that a substantial portion of the gut microbial diversity in sub-Saharan African populations remains poorly characterized, reflecting the limited representation of these populations in current microbiome studies. Consistent with previous studies, taxa such as *Succinivibrio* and *Treponema*, previously associated with African and other non-industrialized populations^17,20^, were also enriched relative to both industrialized and other non-industrialized populations (Supplementary Table 3). In contrast, species specifically depleted in sub-Saharan Africa included species belonging to the genus *Bifidobacterium* (e.g., *B. adolescentis, B. longum*, and *B. bifidum*), as well as *Alistipes putredinis, Parabacteroides merdae*, and *Agathobacter faecis*, many of which are commonly present in industrialized populations.

Because many enriched taxa in sub-Saharan African populations corresponded to poorly characterized or uncultured species, we further examined taxonomic patterns across higher phylogenetic ranks. The analysis revealed that the most strongly enriched taxa in sub-Saharan Africa belonged to the phylum Cyanobacteria, specifically the class Vampirovibrionia (formerly known as Melainabacteria), order *Gastranaerophilales*, the family *Gastranaerophilaceae* (Fig. 2d, Supplementary Table 4). Notably, these represent a non-photosynthetic lineage within Cyanobacteria that is distinct from canonical photosynthetic groups. This strong enrichment at higher taxonomic levels likely reflected the consistent direction of association across multiple *Gastranaerophilales* species, where moderate effect sizes at the species level (Fig. 2b) amplified the signal when aggregated. Significant enrichments were also observed for genera such as *Phascolarctobacterium*_A, *Succinivibrio*, and *Cryptobacteroides*. On the other hand, depleted taxa were dominated by lineages within Actinobacteriota and Bacteroidota, including *Bifidobacterium*, the family *Bifidobacteriaceae*, the order *Actinomycetales*, and the genus *Bacteroides*, consistent with the species-level results.

### Significant enrichment of *Gastranaerophilales* in sub-Saharan African populations

The order *Gastranaerophilales*, identified as the most prominently enriched taxon in the sub-Saharan African group (Fig. 2), has been sporadically detected in the gut environments of humans and other animals; however, its ecological roles and associations with host health remain largely unknown^24,25^, representing a common yet overlooked lineage of the human gut microbiome. To address this, we characterized its global distribution and associations with host and environmental factors.

A comparison of the prevalence of *Gastranaerophilales* across countries revealed that it was detected at the highest frequency in sub-Saharan Africa (79.3%), with particularly high prevalence in countries such as Zimbabwe (97.1%), Burkina Faso (93.9%), and Guinea-Bissau (89.3%) (Fig. 3a, Supplementary Table 5). In contrast, its prevalence was substantially lower in industrialized regions, including East Asia & Pacific (10.5%), North America (19.0%), and Europe & Central Asia (40.2%), although intermediate levels were observed in Latin America & Caribbean (68.0%). These results indicate that *Gastranaerophilales* is not ubiquitously distributed but instead exhibits strong geographic specificity, with a profound enrichment in sub-Saharan Africa.

**Fig. 3.**
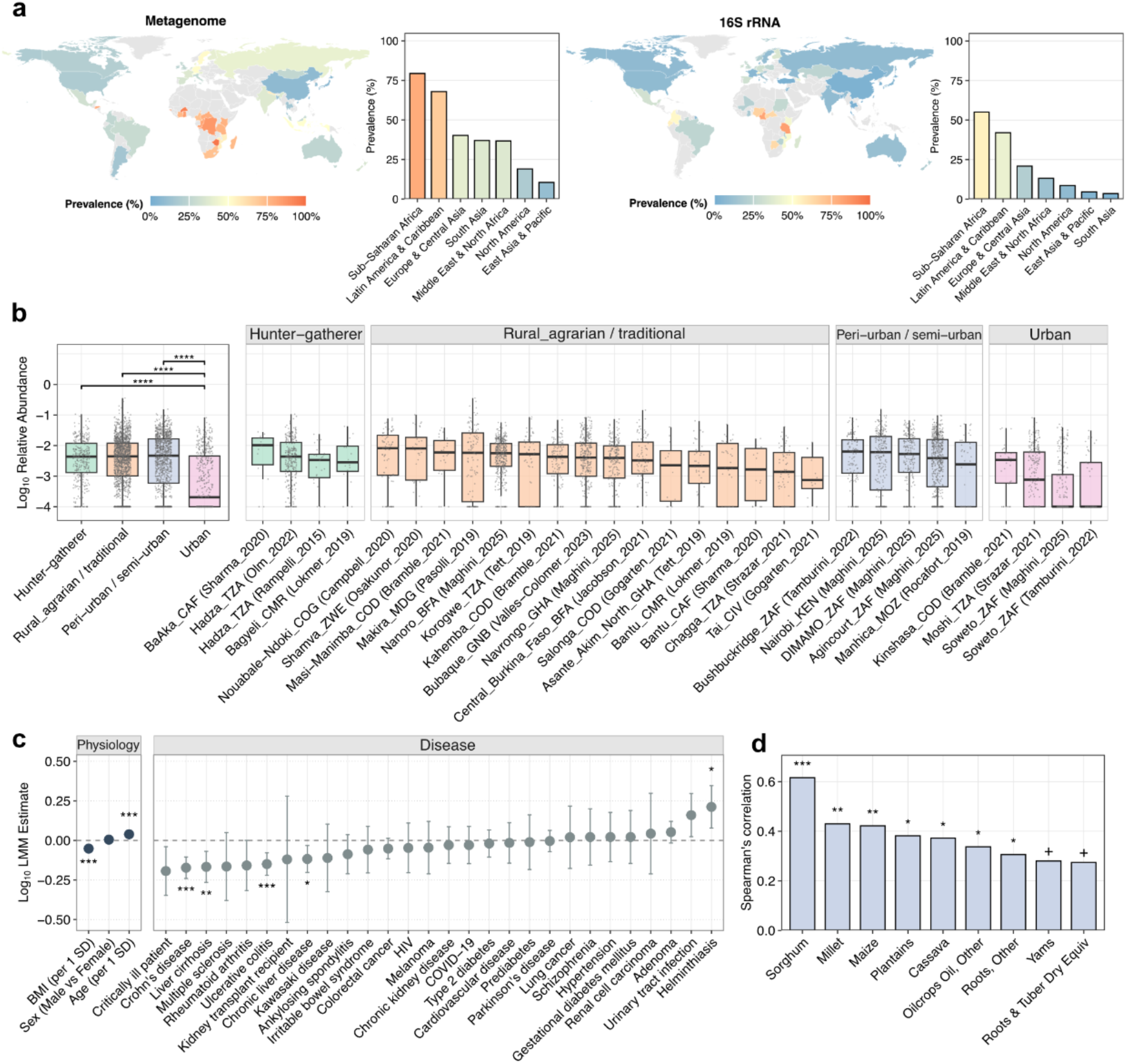
Distribution of the order *Gastranaerophilales* and its association with host factors. **a**, Global prevalence of *Gastranaerophilales* across countries and regions. The left half displays results from metagenomic data (n = 41,471 from 57 countries), and the right half shows 16S rRNA gene data (n = 74,864 from 62 countries). Countries with fewer than 10 samples were excluded from the map. **b**, Variation of *Gastranaerophilales* abundances within sub-Saharan African populations. Samples were categorized into four lifestyle groups (“Hunter-gatherer”, “Rural_agrarian / traditional”, “Peri-urban / semi-urban”, and “Urban”). Box plots show the log_10_-transformed *Gastranaerophilales* relative abundance (with a pseudo-count of 10^−4^) across lifestyles and within specific locations/ethnicities, with individual data points overlaid. The center line indicates the median, box limits indicate the upper and lower quartiles, and whiskers extend to 1.5× the interquartile range. Statistical significance between lifestyle groups was calculated using a two-sided Wilcoxon rank-sum test with Bonferroni correction (****: *P* < 0.0001). **c**, Forest plot showing the association between *Gastranaerophilales* and various host factors. Dots represent LMM estimates, and error bars represent 95% confidence intervals. Asterisks indicate significance levels (*: FDR < 0.05, **: FDR < 0.01, ***: FDR < 0.001). **d**, Spearman’s rank correlation between country-level dietary factors and *Gastranaerophilales* prevalence. Asterisks indicate FDR-adjusted significance levels (+: FDR < 0.1, *: FDR < 0.05, **: FDR < 0.01, ***: FDR < 0.001).

To further disentangle the ecological drivers underlying this distribution, we next focused on within-Africa variation by analyzing the sub-Saharan African samples (n = 3,686). To enable this analysis, we manually curated detailed metadata for each sample, including information on subsistence strategy and degree of urbanization, allowing for a refined classification of lifestyle categories (Methods, Supplementary Table 6). Comparative analysis across the lifestyle categories within sub-Saharan Africa revealed that *Gastranaerophilales* abundance was highest in hunter-gatherers (including the Hadza, BaAka, and Bagyeli), and remained elevated in traditional rural populations primarily engaged in farming and fishing, and transitional rural populations undergoing urbanization (Fig. 3b). In contrast, its abundance was decreased in urban populations. A similar trend was observed for its prevalence, which significantly declined from hunter-gatherer to urban groups (Extended Data Fig. 1). This gradient closely mirrors the degree of urbanization and lifestyle transition, suggesting that *Gastranaerophilales* is associated with traditional ecological contexts and is progressively lost with modernization.

To validate the robustness of the global distribution patterns of *Gastranaerophilales* observed in the metagenomic dataset, we reanalyzed an independent large-scale 16S rRNA dataset from the Human Microbiome Compendium^26^ (Methods). This dataset included 74,864 samples from 62 countries and spanned a broad range of geographic regions, environmental contexts, and population lifestyles, providing a globally diverse dataset for microbiome variation (Supplementary Tables 7 and 8). Despite the known limitations of 16S rRNA-based profiling, including amplification biases and lower taxonomic resolution^27,28^, this dataset also revealed the highest detection rate of *Gastranaerophilales* in sub-Saharan Africa (55.1%) (Fig. 3a, Supplementary Table 9). In regions outside of sub-Saharan Africa, the detection rates were 20.8% for Europe & Central Asia, 8.6% for North America, 4.6% for East Asia & Pacific, and 42.0% for Latin America & Caribbean, closely recapitulating the trends observed in the metagenomic data (Fig. 3a). These results confirm that the significant enrichment of *Gastranaerophilales* in sub-Saharan Africa is robust across independent datasets, sequencing technologies, and population contexts.

### The abundance of *Gastranaerophilales* is associated with host physiology, diseases, and diet

Although *Gastranaerophilales* is detected at a low frequency in populations worldwide, its role in human health and disease remains largely unclear. To investigate its potential biological relevance, we performed an association analysis using LMM incorporating host characteristics (age, sex, and BMI), as well as 32 disease categories, while adjusting for population grouping (i.e, sub-Saharan, other non-industrialized, and industrialized groups). The analysis revealed that the abundance of *Gastranaerophilales* was positively associated with host age and negatively associated with BMI (FDR < 0.05, Fig. 3c). Furthermore, it was significantly enriched in individuals with helminthiasis, whereas it was significantly depleted several chronic inflammatory and metabolic disease status, including inflammatory bowel diseases (Crohn’s disease and ulcerative colitis), and liver cirrhosis (FDR < 0.05, Fig. 3c, Supplementary Table 10). These associations imply that *Gastranaerophilales* may be linked to favorable host physiological states and could represent a potentially beneficial microbial lineage that is reduced under conditions associated with metabolic and inflammatory disease.

Given the strong influence of diet on the composition of the gut microbiome^29,30^, we next examined associations between *Gastranaerophilales* and dietary patterns using country-level food consumption data from the FAOSTAT database (Methods). Correlation analysis based on the metagenomic data revealed significant positive correlations between the prevalence of *Gastranaerophilales* and traditional staple crops characteristic of sub-Saharan African diets, such as sorghum, maize, and millet (Spearman correlations > 0.3, FDR < 0.05, Fig. 3d, Supplementary Table 11). The analysis based on 16S rRNA data also revealed significant positive associations with plantains, starchy roots, and roots & tuber dry equiv (Spearman correlations > 0.3, FDR < 0.05, Extended Data Fig. 2). These crops are rich in dietary fiber and resistant starch, suggesting that *Gastranaerophilales* may occupy a metabolic niche associated with these plant-based, high-fiber diets.

### *Gastranaerophilales* exhibits unique genetic features distinct from other gut bacteria

The genomic characteristics of *Gastranaerophilales* in the human gut were largely unclear. To evaluate its functional potential, we explored representative genomes of the human gut microbiome from the Unified Human Gastrointestinal Genome (UHGG) v2 database^31^ (n = 4,714), including 62 *Gastranaerophilales* genomes (Supplementary Table 12). All *Gastranaerophilales* genomes in UHGG were metagenome-assembled genomes (MAGs), reflecting the absence of cultured representatives. These genomes were almost exclusively assigned to the family *Gastranaerophilaceae* and spanned multiple genera, the majority of which were represented by placeholder annotations (e.g., CAG-, UMGS-, UBA-), with only a limited number of named genera such as *Gastranaerophilus*. High-quality MAGs of *Gastranaerophilales* were relatively compact (1.8–2.3 Mb) and exhibited low GC content (27.4–41.2%).

Principal coordinate analysis based on KEGG Orthology (KO) profiles revealed that *Gastranaerophilales* (Cyanobacteria) formed a unique cluster distinct from other major gut bacterial phyla (Fig. 4a), indicating that its gene repertoire is different from that of other major gut phyla. To identify the functional signatures underlying this distinctiveness, we performed KEGG pathway enrichment analysis. The analysis revealed significant enrichments of several pathways in *Gastranaerophilales*, including flagellar assembly, biosynthesis of cofactors (specifically folate biosynthesis and biotin metabolism), fatty acid biosynthesis, biosynthesis of nucleotide sugars, and lipopolysaccharide biosynthesis (associated with cell wall components) (Fig. 4b, Supplementary Table 13). Notably, while typical photosynthetic cyanobacteria lack flagella, genes related to flagellar assembly were widely conserved in *Gastranaerophilales*. Specifically, approximately 25% of the *Gastranaerophilales* genomes encoded a complete set of flagellar genes, while the remaining retained specific flagellar assembly-related genes (e.g. *flhB, fliI*, and *flgG*), which are postulated to be involved in the extracellular secretion of substances^32^ (Extended Data Fig. 3). This indicates that *Gastranaerophilales*, despite belonging to the phylum Cyanobacteria, had undergone substantial functional divergence from its photosynthetic cyanobacterial relatives and might adapt to the gut environment.

**Fig. 4.**
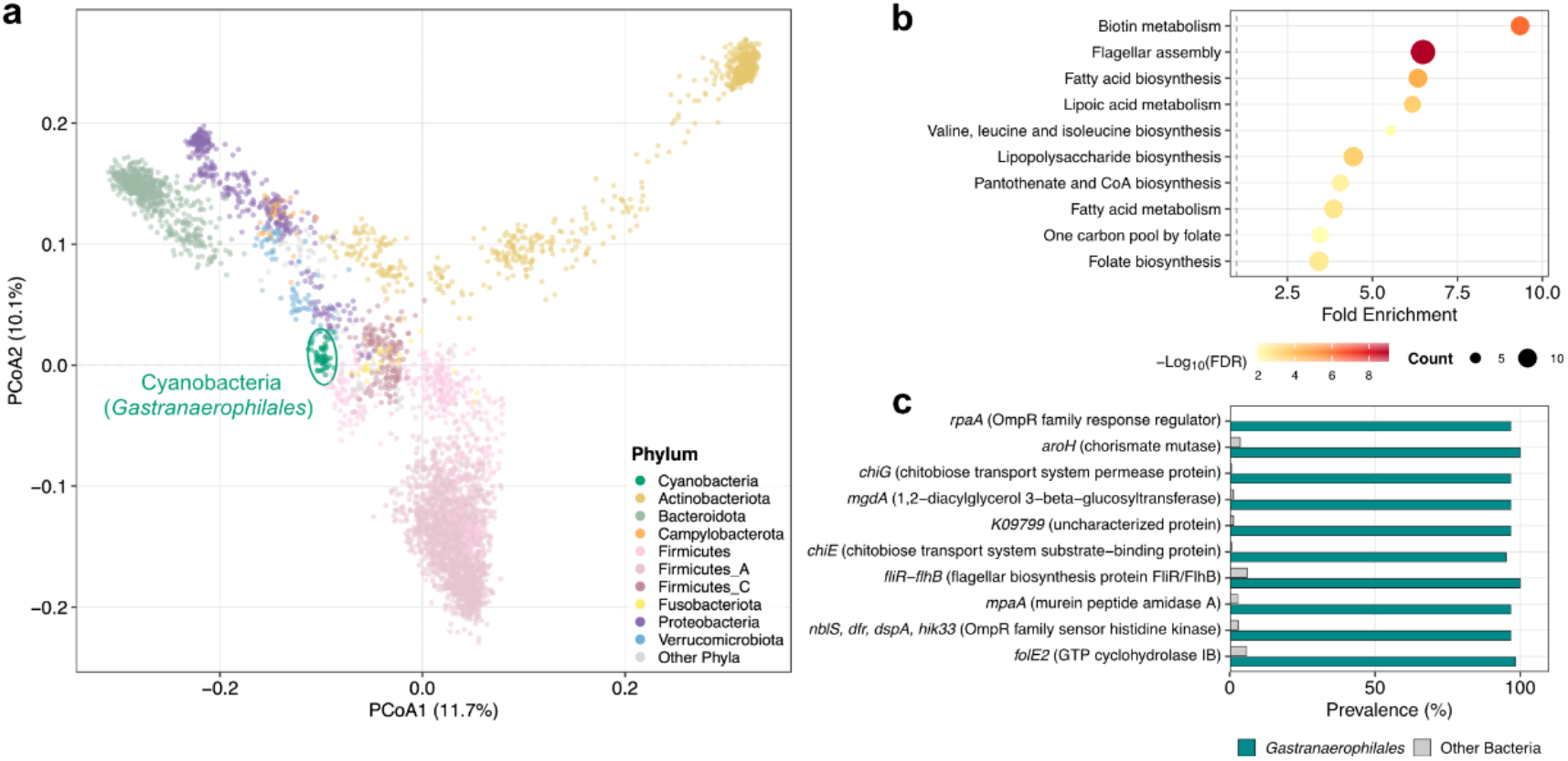
Functional and genomic characterization of human gut *Gastranaerophilales*. **a**, Principal coordinate analysis based on KEGG Orthology profiles of representative genomes from the UHGG database (n = 4,714). Each point represents a bacterial genome. The ten most abundant phyla are shown in distinct colors, whereas all remaining phyla are grouped as other phyla and shown in gray. Cyanobacteria (*Gastranaerophilales*) genomes are highlighted by a green circle. **b**, Enrichment of KEGG pathways in *Gastranaerophilales* compared with other human gut bacteria. The plot shows the top 10 pathways significantly enriched (FDR < 0.05 and prevalence difference > 0.3) in *Gastranaerophilales*, sorted by fold enrichment. Circle size represents the number of genes in the pathway, and color indicates the FDR. **c**, Top KOs specifically enriched in *Gastranaerophilales*. The bar chart displays the prevalence of the top 10 KOs with the largest difference in prevalence between *Gastranaerophilales* and other gut bacterial genomes.

Comparative analysis at individual KO levels further revealed that specific KOs were highly conserved in *Gastranaerophilales* but rare in other human gut species. These included *rpaA* (OmpR family response regulator), *aroH* (chorismate mutase), *chiG* (chitobiose transport system permease protein), *mgdA* (1,2-diacylglycerol 3-beta-glucosyltransferase), and *chiE* (chitobiose transport system substrate-binding protein), all of which were encoded by nearly all *Gastranaerophilales* (>95%), whereas they were rare in other human gut species (<10%) (Supplementary Table 14). Moreover, components of the chitobiose transport system (*chiG, chiE*, and *chiF*: chitobiose transport system permease protein) were among the KOs exhibiting the largest differences in prevalence between *Gastranaerophilales* and other human gut bacterial species (Fig. 4c). These genes encode an ABC transporter responsible for chitobiose, a degradation product of chitin commonly found in insects and fungi.

Taken together, these results indicate that *Gastranaerophilales* possesses a highly specialized functional repertoire that is distinct from other human gut species and may reflect adaptation to ecological niches associated with traditional diets and environmental exposures.

### *Gastranaerophilales* is widely distributed across diverse animal hosts and conserved in primates

To investigate the host range and ecological distribution of *Gastranaerophilales* beyond humans, we additionally collected and analyzed gut microbiome data from 31 non-human animal species (n = 5,371) (Methods, Supplementary Table 15). Across host species, *Gastranaerophilales* exhibited widespread but non-uniform distribution, indicating strong host-associated patterns. High prevalence was observed in several herbivorous and omnivorous mammals, including livestock, particularly *Sus scrofa* (pig, 87%) and *Bos taurus* (cattle, 57%), as well as in avian species such as *Gallus gallus* (chicken, 67%), while moderate detection frequencies were found in ruminants such as *Ovis aries* (sheep, 44%), and *Capra hircus* (goat, 32%) (Fig. 5a). Among Rodents, *Rattus norvegicus* (rat, 89%) exhibited high prevalence, whereas *Mus musculus* (mouse, 24%) showed lower prevalence. In contrast, *Gastranaerophilales* was nearly absent in Carnivorous species, including *Felis catus* (cat, 0%) and *Canis lupus* (dog/wolf, 3%), indicating that its persistence is linked to dietary ecology and gut environment.

**Fig. 5.**
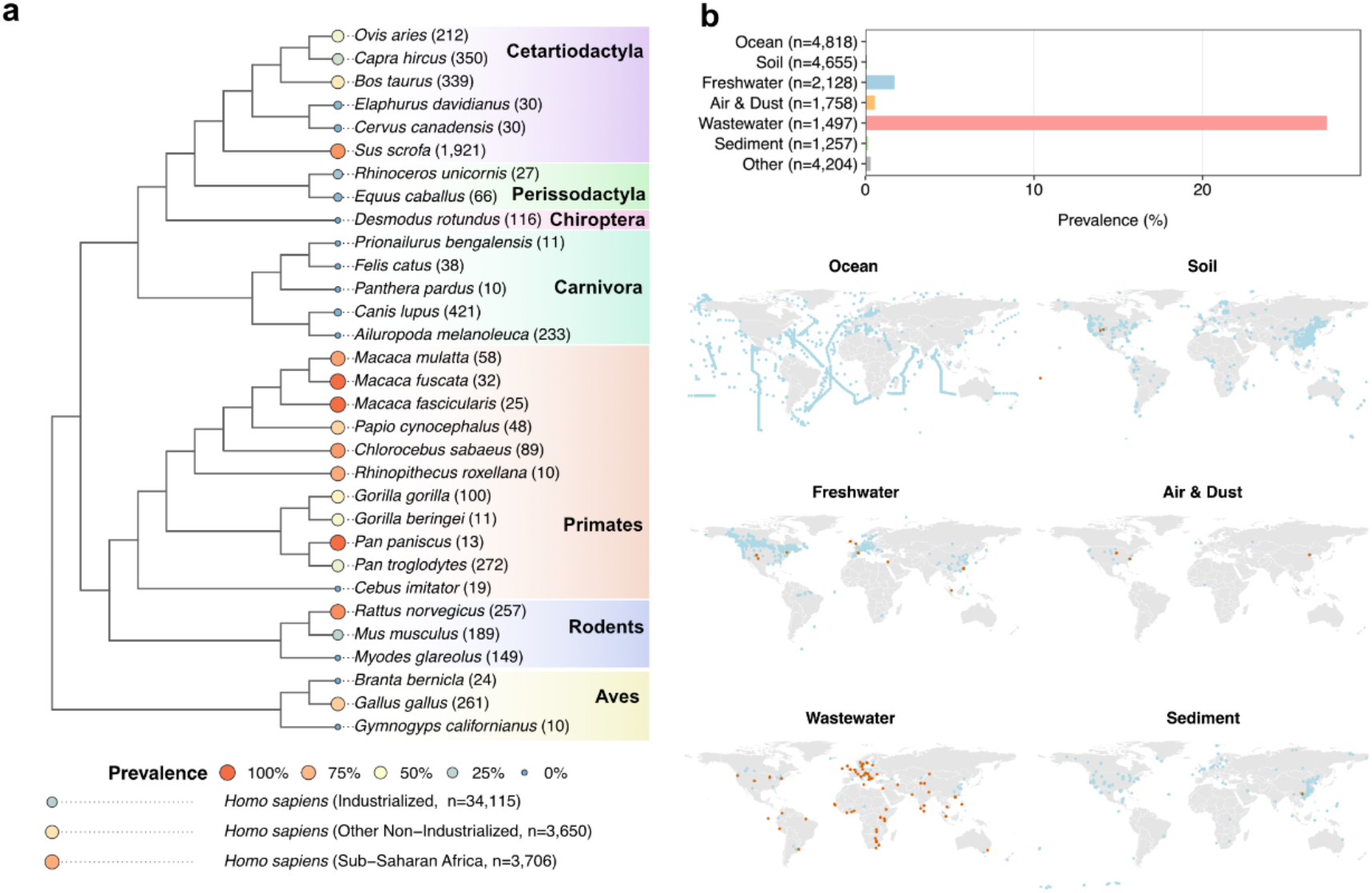
Distribution of *Gastranaerophilales* across non-human animal hosts and environmental ecosystems. **a**, Phylogenetic relationship among diverse animal species and the prevalence of *Gastranaerophilales* in their gut. The dot size and color indicate the prevalence within each animal species. Numbers in parentheses next to the species names indicate the sample size. For comparison, the human prevalence for each analysis group is shown at the bottom. **b**, Detection and frequency of *Gastranaerophilales* across various environmental categories based on global environmental metagenomic data. The bar chart shows the detection frequency within each environmental category. The global map shows the geographic distribution of samples where the taxon was detected (orange) and not detected (blue) for each environment.

Notably, *Gastranaerophilales* was widely conserved across primates, including great apes—our closest evolutionary relatives—including *Pan troglodytes* (chimpanzee, 42%), *Pan paniscus* (bonobo, 100%), *Gorilla beringei* (eastern gorilla, 45%), and *Gorilla gorilla* (western gorilla, 54%). Broad prevalence was also observed across Old World monkeys (catarrhines), including *Macaca mulatta* (rhesus macaque, 83%), *Macaca fuscata* (Japanese macaque, 100%), and *Papio cynocephalus* (yellow baboon, 65%), whereas it was absent in the New World monkey *Cebus imitator* (white-faced capuchin, 0%). The widespread distribution across primates suggests that *Gastranaerophilales* may have established a long-term symbiotic relationship within primate lineages, potentially predating the emergence of modern humans. Similarly, other microbial lineages characteristic of traditional human lifestyles—such as *Prevotella, Succinivibrio*, and *Treponema*—also exhibited broad conservation across non-human primates (Extended Data Fig. 4).

To further investigate its presence in environmental distribution, we analyzed metagenomic datasets from non-host environments (n = 20,317, Supplementary Table 16), including ocean, soil, freshwater, air, wastewater, and sediment samples (Fig. 5b). While *Gastranaerophilales* was detected at relatively high frequency in wastewater (>25%), it was rarely found in other environments such as soil, ocean, and freshwater ecosystems (< 2%). The high detection rate in wastewater likely reflects human and animal fecal influx, whereas its near absence from natural environments indicates that *Gastranaerophilales* is not a free-living taxon but is instead highly adapted specifically to the animal gastrointestinal tracts.

## Discussion

In this study, we systematically characterize the gut microbiome of sub-Saharan African populations using a large-scale microbiome dataset integrating metagenomic (n = 41,471) and 16S rRNA gene sequencing data (n = 74,864). Our analyses revealed that the gut microbiome of sub-Saharan Africans displays distinctive features that clearly differentiate it from both industrialized populations and other non-industrialized groups (Figs. 1 and 2), with a significant enrichment of *Gastranaerophilales*, a lineage of non-photosynthetic cyanobacteria that has received little attention to date (Fig. 3). These findings suggest that dietary or ecological conditions characteristic of sub-Saharan Africa contribute to the persistence of specific microbial lineages, and raise the possibility that potential ancestral microbial taxa largely reduced in industrialized societies remain preserved in African populations.

The order *Gastranaerophilales* belongs to the class Vampirovibrionia (formerly Melainabacteria), a group of non-photosynthetic anaerobic bacteria. To date, no cultured representative of this lineage has been reported, and its evolutionary position has primarily been inferred from phylogenetic analyses of MAGs^24,33,34^. While other orders within Vampirovibrionia (e.g., *Vampirovibrionales, Obscuribacterales, Caenarcaniphilales*) are mainly reported from environmental habitats, *Gastranaerophilales* has been detected in the gastrointestinal tracts of humans and other animals^25^. Despite previous reports of its presence in the human gut at low abundance^33^, its ecological and functional roles have remained largely unexplored. Our comparative genomic analyses revealed that this lineage encodes genes involved in flagellar assembly, as well as biosynthetic pathways for cofactors such as folate and biotin. In addition, *Gastranaerophilales* harbors specialized carbohydrate utilization systems, including transporters for chitobiose (Fig. 4). Consistent with these genomic features, country-level dietary analyses showed positive associations between *Gastranaerophilales* and traditional African staple foods rich in dietary fiber and resistant starch (e.g., sorghum, millet, plantain, and cassava). These results suggest that *Gastranaerophilales* occupy an ecological niche shaped by traditional dietary environments in sub-Saharan Africa.

Analyses of non-human animal gut metagenomes further showed that *Gastranaerophilales* is widely distributed among primates, suggesting that this lineage has established symbiotic associations with the common ancestor of humans and their close relatives (Fig. 5a). Gut bacterial taxa that are common in non-industrialized societies but rare in industrialized populations have been described as VANISH taxa or disappearing microbiota (e.g., *Prevotella, Treponema*, and *Succinivibrio*)^19,20^. Our results suggest that *Gastranaerophilales* also belongs to this group, potentially representing a component of an ancestral gut microbiome shared among humans and related hosts. Notably, even within African populations, individuals living in urban environments exhibited reduced abundances of *Gastranaerophilales* compared with hunter–gatherer or rural populations (Fig. 3b). This pattern suggests that ongoing lifestyle transitions associated with industrialization, including reduced dietary fiber intake, increased consumption of ultra-processed foods, antibiotic exposure, and changes in hygiene environments, contribute to the decline of this lineage in contemporary African populations^35–37^. At the same time, non-communicable diseases such as diabetes and cardiovascular disease are increasing across Africa, and alterations in gut microbial ecosystems have been proposed as a contributing factor^38,39^. Further studies will therefore be important to clarify the functional roles of *Gastranaerophilales* in the human gut and its association with host physiology and disease.

In summary, our study demonstrates that sub-Saharan African populations harbor gut microbiome structures that differ markedly from those observed in industrialized societies. In particular, the enrichment of the previously underexplored lineage *Gastranaerophilales* suggests the regional preservation of an ancestral symbiotic lineage in Africa. Our findings underscore the importance of studying the human gut microbiome across the full diversity of human populations, cultures, and environments. Integrating this diversity will be essential for reconstructing the evolutionary history of the human–microbe relationship and for developing a more complete understanding of the global diversity of the human gut microbiome.

## Methods

### Human gut microbiome dataset

To analyze the global diversity of the human gut microbiome, we utilized Metalog^22^, a large-scale repository of public metagenomes. Species-level taxonomic profiles of the human gut microbiomes and associated metadata (age, sex, BMI, disease, country, and study) were retrieved from this database (accessed in January 2026). Samples from infants (under 3 years old), whose gut microbiome composition significantly differs from that of older children and adults^10^, as well as samples from individuals treated with antibiotics or fecal microbiota transplantation, were excluded from the analysis. Consequently, a total of 39,652 samples from 56 countries were obtained. Furthermore, we integrated 1,819 samples from AWI-Gen 2 Microbiome Project^21^— currently the largest gut microbiome study in Africa. These samples were analyzed using mOTUs v3.0.0, consistent with the Metalog dataset. The combined dataset comprises 41,471 samples from 57 countries worldwide. In addition to the taxonomic information provided by mOTUs, those based on the Genome Taxonomy Database (GTDB)^23^ were assigned to each bacterial species based on the conversion table provided by mOTUs. To analyze microbiome profiles at higher taxonomic ranks, species-level relative abundances were aggregated to the phylum, class, order, family, and genus levels based on the GTDB taxonomy.

For independent validation, a 16S rRNA gene amplicon sequencing dataset was also collected from the Human Microbiome Compendium (https://zenodo.org/records/13733642)^26^. To ensure data quality, metadata were manually curated. Samples from non-human animal models and infants, data with an average read length of less than 100 bp, and data lacking information on the country of sample collection were excluded. Data from the United States Minor Outlying Islands were integrated to those from the United States. The final 16S rRNA dataset comprised 74,864 samples from 62 countries.

### Statistical analysis

Data formatting, statistical analysis, and visualization in this study were performed using R (version 4.3.3)^40^. Data manipulation and visualization were primarily conducted using the tidyverse package suite (version 2.0.0)^41^, the patchwork package (version 1.3.1)^42^, and the ggpubr package (version 0.6.1)^43^. Map rendering and spatial data retrieval were performed using the rnaturalearth (version 1.0.1)^44^ and sf (version 1.0.21)^45^ packages.

### Microbiome-based population grouping and differential abundance analysis

Principal component analysis (PCA) was performed on the species-level taxonomic profiles using the prcomp function. To reduce analytical noise, only major species with an average relative abundance of ≥0.01% and a prevalence of ≥10% were included in the analysis. Prior to PCA, a pseudo-count of 10^−4^ was added to the profile, and a log_10_ transformation was performed. Countries were grouped into clusters of industrialized and non-industrialized groups, based on the PC2 score of Malaysia, which was located at the boundary of the clusters (Fig. 1b). Furthermore, considering geographical factors, the non-industrialized group was divided into the sub-Saharan African group and the remaining non-industrialized group. Subsequent comparative analyses were conducted among these three groups.

Alpha diversity of the microbiome was assessed using the Shannon index (diversity function of the vegan package version 2.7-1)^46^, and observed species richness (number of species with a relative abundance of ≥0.01%). Pairwise comparisons among the three groups were performed using the Wilcoxon rank-sum test, followed by multiple testing correction via the Bonferroni method.

To test for differences in the relative abundance of microbial species among the three groups, a linear mixed model (LMM) was employed. Species with an average relative abundance of ≥0.01% and a prevalence of ≥10% in at least one of the three groups were included (n = 1,064). The response variable was the log_10_-transformed relative abundance of each species with a pseudo-count of 10^−4^. The analysis group (sub-Saharan Africa, other non-industrialized, and industrialized) and disease status (major diseases with ≥ 100 samples and healthy controls) were included as fixed effects, while the country and study were incorporated as random effects. The models were constructed using the lmer function of the lme4 (version 1.1.37)^47^, and p-values were obtained using lmerTest (version 3.1.3)^48^ packages. Estimated marginal means for each group and pairwise comparisons were calculated using the emmeans package (version 1.11.1)^49^ with the Satterthwaite method.

To identify species specifically associated with sub-Saharan Africa, a dual-filtering approach was applied. Species were defined as significantly enriched or depleted only if they met all of the following criteria in both comparisons (sub-Saharan Africa vs. industrialized and other non-industrialized): (1) a false discovery rate (FDR) < 0.05 (Benjamini-Hochberg [BH] method), (2) absolute effect size > 0.1, and (3) consistent direction of change (either enriched or depleted). For ranking and visualization, the more conservative (i.e., smaller absolute) estimate from the two comparisons was adopted as the representative effect size.

Phylogenetic characteristics were further evaluated across taxonomic ranks (phylum to genus). Using LMMs with the same filtering and covariate adjustment. The numbers of taxa retained after filtering were as follows: phylum: 20/45, class: 25/93, order: 54/214, family: 107/489, and genus: 449/2426.

A phylogenetic tree was obtained from GTDB (Release 207). Taxa that could be matched between GTDB and mOTU were retained, and unmatched taxa were excluded from the phylogenetic tree. Tree visualization and annotation were performed using the ggtree (version 3.10.1)^50^ and ggtreeExtra packages (version 1.12.0)^51^.

### Distribution of *Gastranaerophilales* and association with host factors

To investigate the geographical distribution of *Gastranaerophilales*, we calculated its prevalence by country and region for both the metagenomic and 16S rRNA gene datasets. A sample was considered positive if the relative abundance of *Gastranaerophilales* exceeded 0.01%. To ensure robustness, countries with fewer than 10 samples were excluded from the analysis.

To explore variation of *Gastranaerophilales* in the sub-Saharan African populations, we manually collected and curated detailed sample metadata beyond those available in Metalog from the original publications (e.g., region of collection, ethnicity, and study background). Based on the information, samples from sub-Saharan Africa (n = 3,686) were classified into four lifestyle categories: “Hunter-gatherer”, “Rural agrarian/traditional”, “Peri-urban/semi-urban”, and “Urban”. These lifestyle categories were based on previously established classifications^21,52^, with minor modifications to accommodate the present dataset. Differences in the relative abundance of *Gastranaerophilales* among these categories were assessed using a pairwise Wilcoxon rank-sum test with Bonferroni correction for multiple testing.

To assess associations between *Gastranaerophilales* and diseases or host physiological factors, LMM was applied. The response variable in the models was the log_10_-transformed relative abundance of *Gastranaerophilales*, with a pseudo-count of 10^−4^ added prior to transformation. To minimize sample loss due to missing metadata, two separate LMMs were constructed: a disease model and a physiological factor model. In both models, country and study were included as random effects, and the analysis group (sub-Saharan Africa, other non-industrialized, industrialized) was included as a fixed-effect covariate. In the disease model, disease groups with fewer than 100 samples were integrated into an “Other” category and included as a fixed effect, with the control group as the reference. The physiological factor model included samples with complete information on age, sex, and Body Mass Index (BMI) (n = 11,480), with age and BMI standardized as Z-scores, and sex incorporated as fixed effects. Model estimates and 95% confidence intervals were extracted, and the FDR was calculated using the BH method.

To evaluate associations between diets and *Gastranaerophilales*, national food consumption data (supply per capita per day [g/capita/day]) were obtained from the FAOSTAT database (https://www.fao.org/faostat/, accessed November 2024). Spearman’s rank correlation coefficients were calculated between the prevalence of *Gastranaerophilales* and the consumption of each food group (10-year average from 2012 to 2021) in each country using the corr.test function of the psych package (version 2.5.6)^53^. P-values were adjusted using the BH method. To ensure reliability, countries with a sample size of ≤ 20 in the 16S rRNA dataset were excluded.

### Analysis of gene functions and metabolic pathways

To characterize the functional potential of *Gastranaerophilales*, the Unified Human Gastrointestinal Genome (UHGG) database (version 2.0.2)^31^ was employed. In this analysis, only bacterial genomes were included, and archaeal genomes were excluded. Gene annotations by eggNOG-mapper^54,55^, was retrieved, and the presence/absence of KEGG (Kyoto Encyclopedia of Genes and Genomes) Orthology (KO)^56,57^ terms was tabulated for the 4,714 bacterial representative genomes. Functional profiles were compared using principal coordinate analysis (PCoA) based on Jaccard distances calculated from KO presence/absence matrices, using the cmdscale function. To identify KOs characteristic of *Gastranaerophilales*, KO prevalence was compared between the genomes of *Gastranaerophilales* and those of the other taxonomies, using the Wilcoxon rank-sum test with multiple testing correction by the BH method. KOs with FDR < 0.05 and a difference in prevalence > 0.3 were defined as significantly enriched. Pathway enrichment analysis was conducted using the enricher function of the clusterProfiler (version 4.10.1)^58^ and KEGGREST (version 1.42.0)^59^ packages, based on the hypergeometric test. Fold enrichment was calculated as (k/n)/(M/N), where k is the number of significant KOs in the pathway, n is the total number of significant KOs, M is the number of background KOs in the pathway, and N is the total number of background KOs. Pathways with fewer than 5 or more than 500 genes were excluded from the analysis.

### Global distribution of *Gastranaerophilales* across animal hosts and environmental ecosystems

To investigate the distribution of *Gastranaerophilales* outside the human gut, we obtained species-level taxonomic profiles of gut metagenomes from non-human animals from the Metalog database (n = 5,371 samples from 31 different species). To ensure analytical robustness, subspecies-level entries were integrated at the species level, and only species with ≥10 samples were included in the analysis. As in the human gut metagenomic dataset, GTDB-based taxonomic information was assigned to each microbial species using mOTU IDs. For each animal species, prevalence was defined as the proportion of samples in which the relative abundance of *Gastranaerophilales* exceeded 0.01%. A phylogenetic tree^60,61^ representing the target animal species was retrieved from the Open Tree of Life project using the rotl package (version 3.1.0)^62^ using the ape package (version 5.8.1)^63^ and visualized using the ggtree package.

To evaluate the distribution of *Gastranaerophilales* in non-host environments, species-level taxonomic profiles of environment-derived metagenomes were obtained from the Metalog database (n = 20,317). As in the animal dataset, GTDB-based taxonomic information was assigned to each bacterial species using mOTU IDs. Samples were classified into major environmental categories (i.e., marine, soil, freshwater, wastewater, air/dust, and sediment) based on the environmental information included in the metadata table from Metalog. Samples not fitting into these major categories were treated as “Other”. For each category, samples with a relative abundance of *Gastranaerophilales* exceeding 0.01% were considered positive, and detection frequencies were calculated. To visualize the geographical distribution of *Gastranaerophilales*-positive samples, samples were plotted on a global map using latitude and longitude information from the metadata table from Metalog. For mapping, samples lacking coordinate information or containing invalid coordinate values (e.g., latitude outside ±90° or longitude outside ±180°) were excluded.

## Data availability

The species-level taxonomic profiles of human gut metagenomic samples (n = 39,652 samples from 56 countries), as well as animal and environmental samples (n = 25,688) used in this study are available through the Metalog database (https://metalog.embl.de/). Additional metagenomic samples from the AWI-Gen2 project analyzed in this study (n = 1,819 samples) are available under the accession number PRJNA115737 at the NCBI Sequence Read Archive. The 16S rRNA gene dataset from the Human Microbiome Compendium (n = 74,864 samples from 62 countries) is available at Zenodo (https://zenodo.org/records/13733642).

## Code availability

All custom scripts used for data processing, statistical analyses, and figure generation are publicly available at https://github.com/Kotaro-Murai/GutMicrobiome_Africa_2026.

## Acknowledgments

We thank Michael Kuhn and Anthony Fullam (European Molecular Biology Laboratory) for their valuable assistance with data collection from Metalog. We also thank members of the Suzuki lab for helpful discussions and support with data analysis throughout this study. This work was supported in part by JSPS KAKENHI (Grant Number JP25K23685), the Mitsubishi Foundation, the Takeda Science Foundation, and the AMED PRIME program of the Japan Agency for Medical Research and Development.

