## Extended_Data_Figures for "Global-scale microbiome analyses identify an ancestral gut cyanobacterial lineage enriched in African populations"

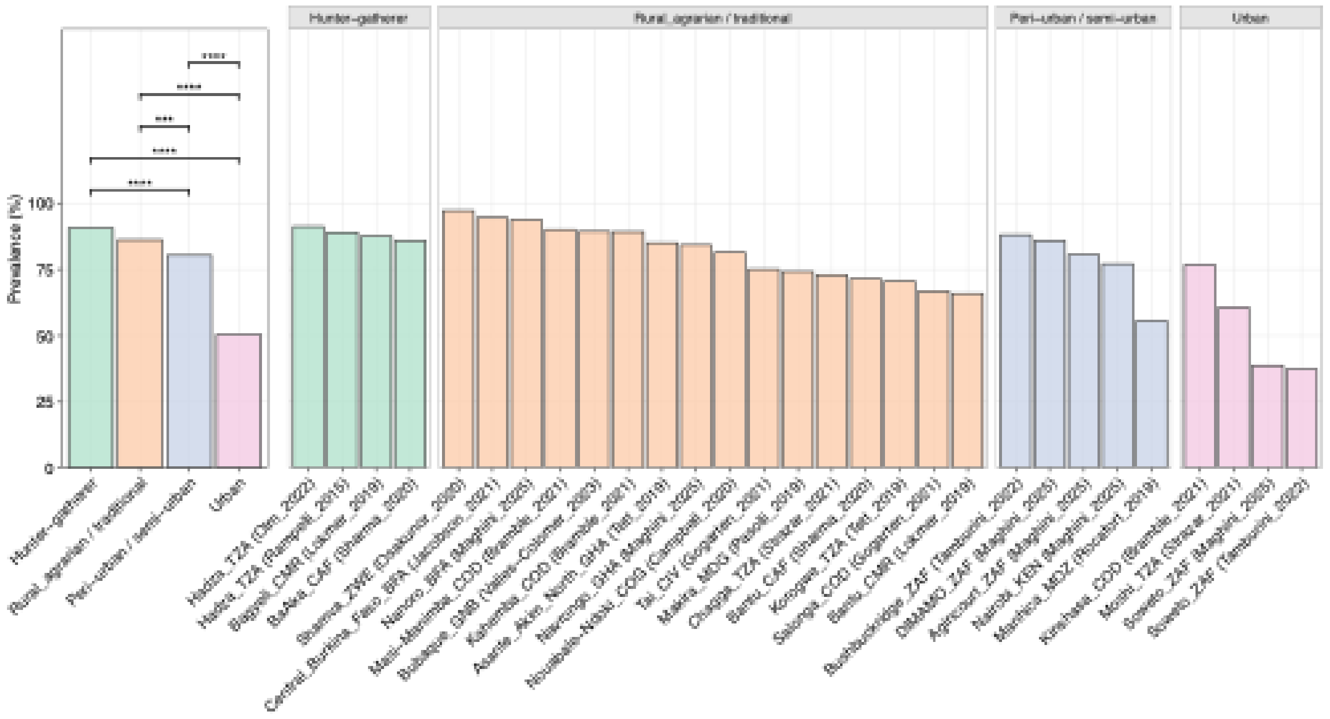


**Extended Data Fig. 1 | Variation in the prevalence of Gastranaerophilales within sub-Saharan African populations.**
Bar charts display the prevalence of the order *Gastranaerophilales* (defined as the proportion of samples with a relative abundance > 0.01%) within sub-Saharan African populations aggregated by lifestyle (left) and stratified by sampling location and study (right). Pairwise comparisons between lifestyle groups were performed using two-sided Fisher's exact tests with Bonferroni correction. Asterisks indicate adjusted P values (*: P < 0.05, **: P < 0.01, ***: P < 0.001, ****: P < 0.0001).

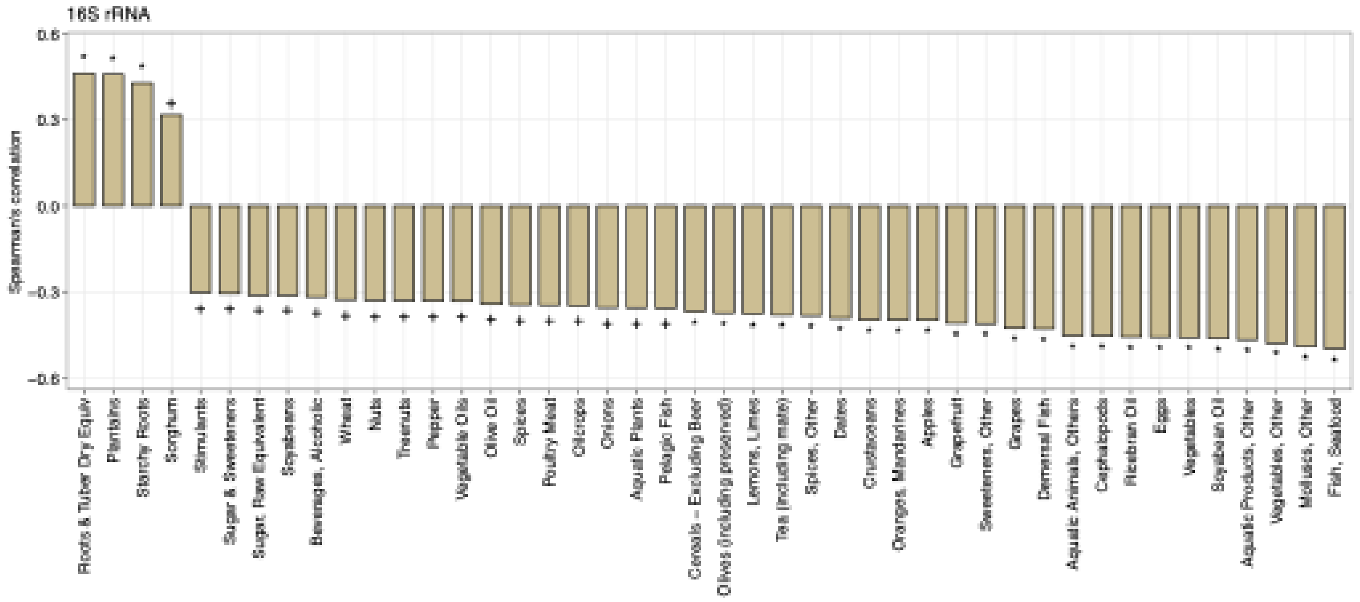


**Extended Data Fig. 2 | Correlations between diets and *Gastranaerophilales* in the 16S rRNA dataset.**
Spearman's rank correlation between country-level dietary factors in the FAOSTAT database and *Gastranaerophilales* prevalence. Asterisks indicate FDR-adjusted significance levels (+: FDR < 0.1, *: FDR < 0.05, **: FDR < 0.01, ***: FDR < 0.001). Only dietary factors with an FDR < 0.1 are shown.

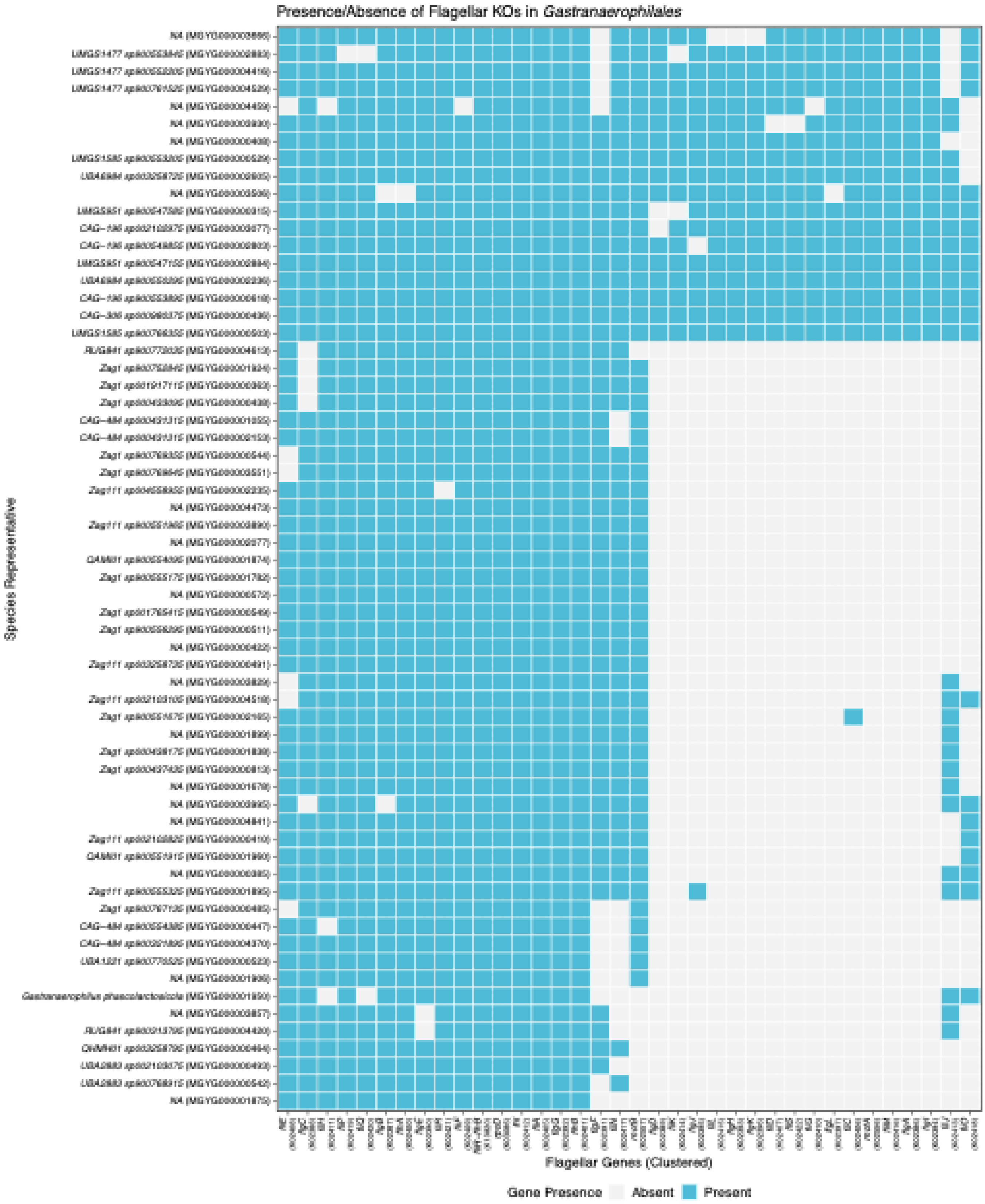


**Extended Data Fig. 3 | Presence and absence of flagellar assembly genes in *Gastranaerophilales*.**
Clustered heatmap displaying the distribution of flagellar assembly genes (KEGG pathway map02040) across *Gastranaerophilales* species representatives from the UHGG database. Rows (genomes) and columns (genes) are ordered by hierarchical clustering using Jaccard distance. Blue tiles indicate gene presence; gray indicates absence.

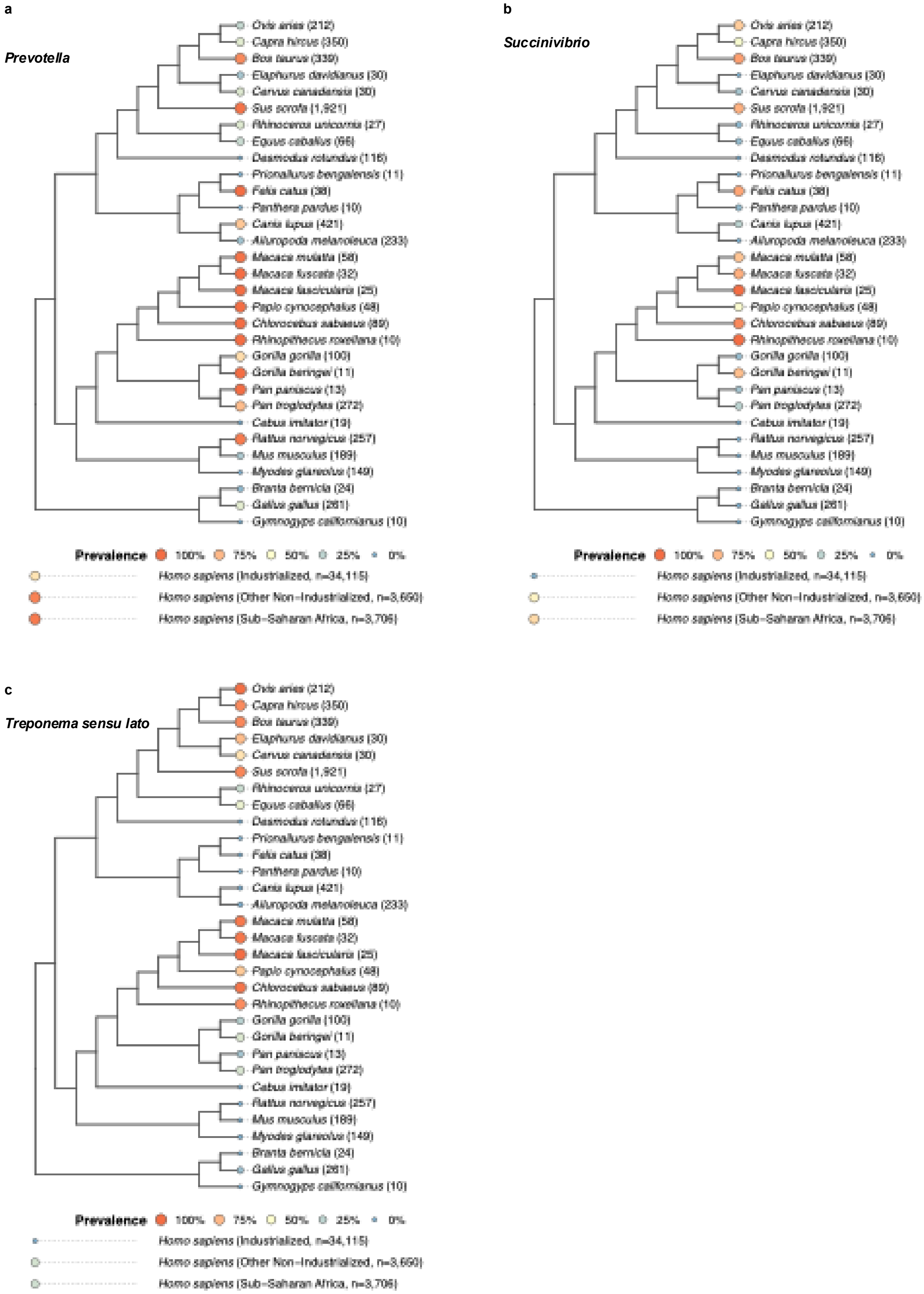


**Extended Data Fig. 4 | Phylogenetic distribution of *Prevotella*, *Succinivibrio*, and *Treponema sensu lato*.**
Prevalence of **a,** *Prevotella*, **b,** *Succinivibrio*, and **c,** *Treponema sensu lato* mapped onto animal phylogenies and human cohorts categorized by lifestyle. Cirde size and color denote the prevalence within each host species or human group. Detection was defined as a relative abundance > 0.01%. Numbers in parentheses next to the species names indicate the sample size. For *Treponema sensu lato*, the relative abundance represents the sum of all GTDB-defined *Treponema* lineages (including *Treponema_A*, *Treponema_B*, etc.). Only animal species with ≥10 samples are included.
